# Unveiling the Power of PROTAC Valency: Navigating Cell Type-Specific Hook Effects

**DOI:** 10.1101/2024.06.12.598404

**Authors:** Arash Chitsazan, Frederik Eisele, Parisa Rabieifar, Hyunsoo Park, Markus Nordberg, Jianming Liu, Stefan Geschwindner, Göran Dahl

## Abstract

Targeted protein degradation (TPD) using bivalent proteolysis-targeting chimeras (PROTAC) technology has shown potential in expanding the “druggable” proteome. In their publication in *Nature Chemical Biology*, Imaide et al.^1^ posited that augmenting PROTAC valency could potentially lead to the formation of long-lived ternary complexes between PROTAC, the protein of interest (POI), and E3 ligase, thereby constraining the formation of potent binary complexes, as evidenced by a pronounced hook effect. The authors introduced SIM1, a trivalent von Hippel–Lindau (VHL)-based PROTAC, which exhibited a superior degradation profile in comparison to its parent molecule MZ1, towards bromo and extra terminal (BET) proteins, with a predilection for BRD2. The authors attributed this heightened degradation capability of SIM1 over bivalent MZ1 as supportive evidence for their hypothesis. While we concur with the notion that increasing valency and avidity could enhance the efficacy of a PROTAC, the claim that trivalent PROTACs unequivocally eliminate the hook effect is not entirely accurate. We propose that the presence or absence of a hook effect is influenced by numerous factors beyond PROTAC valency.

## Methods and Results

Several data from Imaide et al.^1^ prompt consideration of potential differences in the behavior of HiBiT variants of BET proteins compared to the non-modified endogenous proteins in naïve HEK293 cells. In naïve HEK293 cells, SIM1 preferred the degradation of BRD4 over BRD2 (fig. 1e), while in CRISPR/Cas9 engineered HEK293 cell lines, for endogenous tagging of HiBiT-BRD2 and HiBiT-BRD4 (HiBiT-BRD2 and HiBiT-BRD4 cells), HiBiT-BRD2 degradation was preferred (fig. 2a). It is noteworthy that SIM1 induced the degradation of HiBiT-BRD2 and HiBiT-BRD4 with a DC_50_ of 60 and 100 picomolar (pM) respectively (fig. 2b), while in naïve HEK293 cells, the efficacy of SIM1 was substantially lower, reported at 1100 and 700 pM for BRD2 and BRD4, respectively (fig. 1e).

**Fig. 1.**
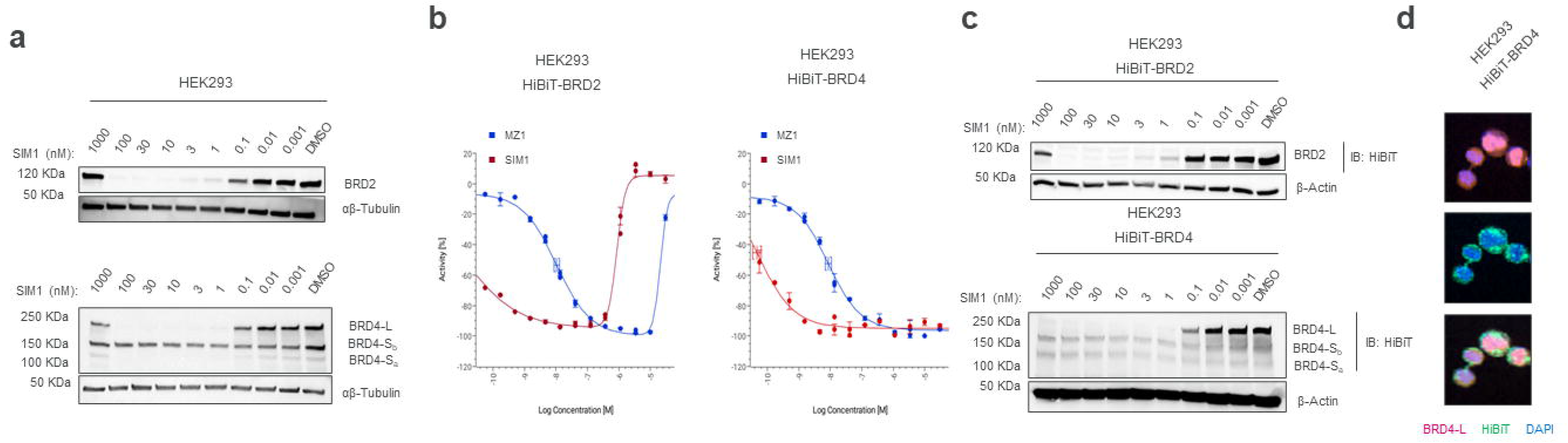
**a**, Immunoblot analysis of BRD2, and BRD4 isoforms after 4 h treatment of naïve HEK293 cells with 9 concentrations of SIM1 or DMSO (n=1). **b**, Quantitative analysis of loss of luminescence due to degradation HiBiT-tagged BET protein using 13-point dose response for HiBiT-BRD2 and HiBiT-BRD4 in HEK293 cells following 4 h treatment with MZ1, SIM1 or DMSO (3 technical replicates). **c**, Immunoblot analysis of HiBiT-BRD2, and HiBiT-BRD4 isoforms in HEK293 HiBiT-BRD2 and HEK293 HiBiT-BRD4 cells with 9 concentrations of SIM1 or DMSO (n=2). **d**, Immunofluorescence of BRD4-L and total HiBiT in HEK293 HiBiT-BRD4 cells cultured overnight in full media. Nuclei stained with DAPI (scale bar, 20 μm).

**Fig 2.**
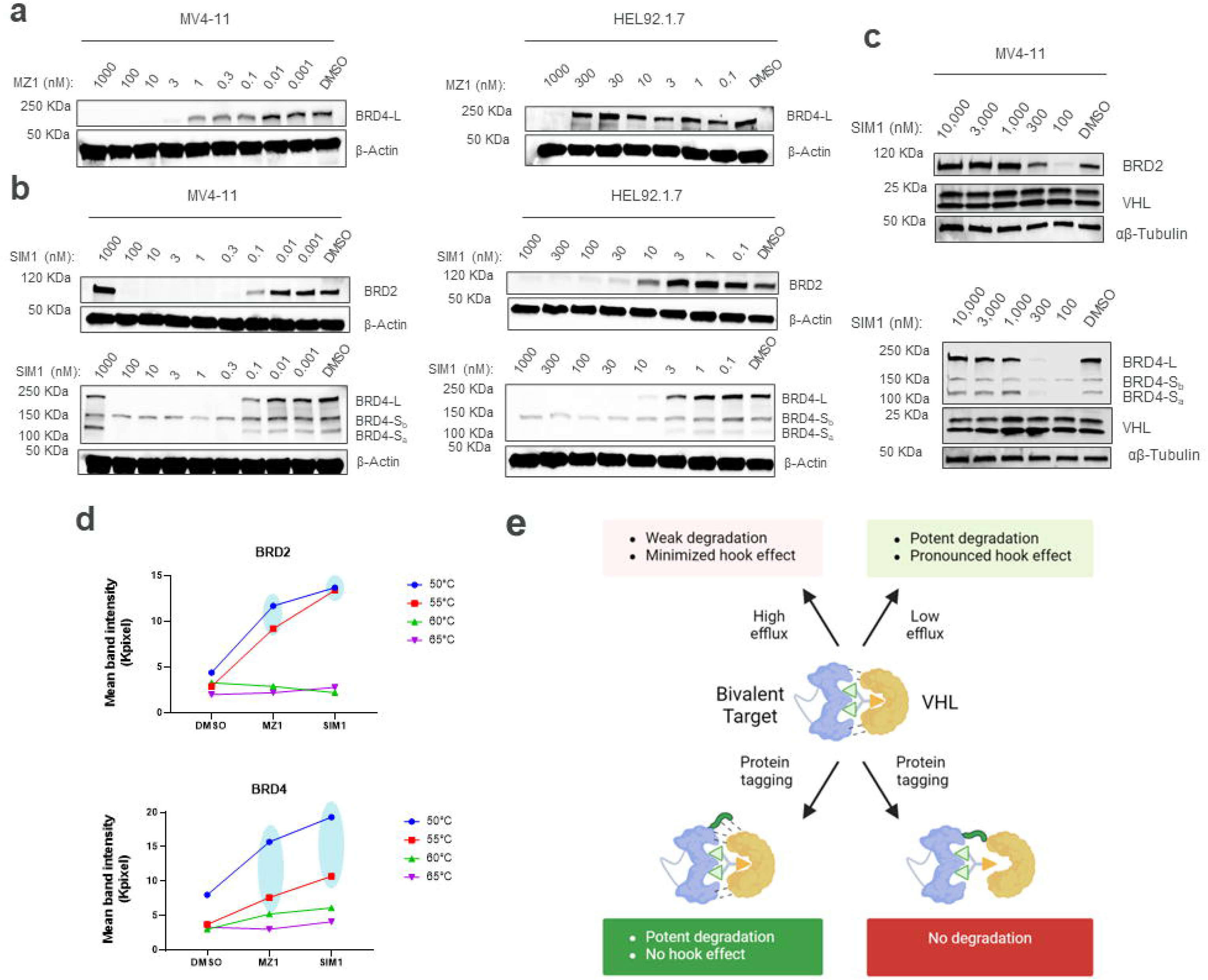
**a**, Immunoblot analysis of BRD4-L after 4 h treatment of MV4-11 cells (left) and HEL92.1.7 cells (right) with MZ1 (1 pM–1000 nM or 100 pM–1000 nM for MV4-11 and HEL92.1.7 cells respectively) or DMSO. **b**, Immunoblot analysis of VHL, BRD2 and BRD4 isoforms after 4 h treatment of MV4-11 cells (left) and HEL92.1.7 cells (right) with SIM1 (1 pM–1000 nM or 100 pM–1000 nM for MV4-11 and HEL92.1.7 cells respectively) or DMSO. **c**, Immunoblot analysis hook effect for BRD2 and BRD4 isoforms after 4 h treatment of MV4-11 cells with SIM1 (100 nM–10 μM). **d**, Quantitative analysis of lysate-CETSA experiment conducted at 50-65°C (5°C interval) lysate of naïve HEK293 cells. Immunoblot performed from remaining soluble BRD2 and BRD4 proteins following 20 minutes incubation of 100 μM MZ1 or SIM1 with the lysate of naïve HEK293 cells at room temperature (n=1). **e**, Cartoon depicting the impact of PROTAC efflux, favourable and unfavourable genetic modifications of POI on TPD.

Hence, we compared the degradation profile of SIM1 in naïve HEK293 cells and the HiBiT-BRD2, and HiBiT-BRD4 cells (Fig. 1). SIM1 degraded both the BRD4-L and BRD4-Sa isoforms, but not BRD4-Sb in naïve HEK293 cells (Fig. 1a). A pronounced hook effect was detected for BRD2 and, to a lesser extent, for BRD4 in naïve HEK293 cells (Fig. 1a). This finding significantly differs from the results presented by Imaide et al., where no hook effect was observed for SIM1 at 1μM (fig. 1d and e). We further characterized the hook effect in HiBiT-BRD2 and HiBiT-BRD4 cells, using Nano-Glo HiBiT lytic detection assays. This confirmed the presence of a hook effect for HiBiT-BRD2 for both SIM1 and MZ1, but no hook effect was detected for HiBiT-BRD4, even at high concentrations (30 μM) (Fig. 1b). Additionally, in contrast to the results presented by Imaide et al., we did not observe a significant difference in the efficiency with which SIM1 degraded non-tagged and HiBiT-tagged BRD2 and BRD4.

Subsequently, we utilized HEK293 cells that stably express LgBiT and HiBiT-BRD4 to investigate even higher concentrations of SIM1. Notably, no hook effect was detected, even at high micromolar concentrations of SIM1 (30 μM) (Supplementary Fig. 1a). However, both MZ1 and SIM1 exhibited slightly lower efficacy in degrading the larger LgBit:HiBiT-BRD4 complex compared to the smaller HiBiT-BRD4 protein (Supplementary Fig. 1a and Fig. 1b, respectively). We further investigated the SIM1 mediated degradation profiles in HiBiT-BRD2/BRD4 cell lines by Western blot against HiBiT. The result confirmed the hook effect with SIM1 in HiBit-BRD2 cells and no hook effect in HiBiT-BRD4 cells for HiBit-BRD4-L (Fig. 1c).

While the degradation of BRD4-L appeared similar as in naïve HEK293 cells, no degradation of the short BRD4 isoform (HiBiT-BRD4-Sa) was detected in the HiBiT-BRD4 cells. Moreover, dual immunostaining of HiBit-BRD4 cells against the C-terminus of BRD4-L and HiBiT revealed a different nuclear localization of total HiBiT-BRD4 isoforms from the HiBit-BRD4-L (Fig. 1d). This indicates that the SIM1 degradation efficacy for tagged versus non-tagged BRD4 could be different in sub-cellular localization, something that has also been described by others^2^ (Simpson et al. 2022).

We proceeded to investigate the degradation profile of SIM1 in other cell lines expressing non-modified BRD2 and BRD4. In their fig 1f, Imaide et al. utilized MV4-11 acute leukemia cells for a cell viability test. We observed a significant discrepancy between the reported IC_50_ for SIM1 and MZ1 (fig. 1f) in MV4-11 cells and the DC_50_ in HEK293 cells. We hypothesized that this variation might be attributed to differing levels of MultiDrug Resistance protein 1 (MDR1)-mediated efflux of PROTACs, as reported for the parental MZ1^3^ (Kurimchak et al. 2023).

To delve deeper into this hypothesis, we compared the mRNA expression profile of *ABCB1* (which encodes MDR1) in MV4-11 cells (DepMap portal)^4^ to a panel of other suspension cells and observed that MV4-11 cells exhibit very low ABCB1, while the expression levels of *ACTB, BRD4*, and *VHL* were comparable to those of other cells (Supplementary Fig. 1b). Among others, HEL92.1.7 cells, which showed high MDR1 expression, was chosen as a high efflux cell model in our study. The translation of mRNA level to protein level in different cell lines was confirmed by Western blot (Supplementary Fig. 1c.)

In low efflux MV4-11 cells, MZ1 showed potent degradation of BRD4-L following 4 hours of incubation (D_max_ at 3,000 pM) without a hook effect(Fig. 2a) while SIM1 treatment improved the efficacy but led to a pronounced, and weak hook effect for BRD2 (D_max_ at 300 pM) and BRD4 (D_max_ at 300 pM) respectively (Fig 2b). Since MDR1-mediated efflux does not play a significant role in MV4-11 cells, the 10-fold increase in efficacy between SIM1 and MZ1 can be attributed to the increased PROTAC valency for SIM1. In the high efflux HEL92.1.7 cells, on the other hand, both MZ1 and SIM1 were about 300-fold weaker in degrading BRD4-L, (Fig. 2a and 2b), and SIM1 was about 300-fold weaker in degrading BRD2 (Fig. 2b). Intriguingly, this means that the bivalent MZ1 is a better degrader of BRD4 in low efflux MV4-11 cells than the trivalent SIM1 is in high efflux HEL92.1.7 cells. Within a cell line however, the degradation efficiency difference between MZ1 and SIM1 remains the same. While MDR1 expression is an important differentiating factor between MV4-11 and HEL92.1.7 cells, it is not possible to rule out the importance of other factors that could potentially impact the efficacy of the PROTACs. BRD4-L degradation profiles of both MZ1 and SIM1 being perturbed equally when going from MV4-11 to HEL92.1.7 cells implies that a shift in the intracellular PROTAC concentration is a plausible explanation. While the potency shift for SIM1 and MZ1 was similar, this may not always be the case as different PROTACs could be better or worse substrate for MDR1. Hence, it is crucial to consider intrinsic cellular differences while conducting PROTAC profiling or cell viability assays and use proper controls to assess what drives PROTAC potency.

Next, we employed MV4-11 cells to explore the onset of the hook effect of SIM1 and any potential impact of VHL levels. Using higher concentrations of SIM1, we observed hook effect for BRD2 at 300 nM and for BRD4 at 1000 nM, while VHL protein levels remained stable (Fig. 2c). Therefore, the presence of the hook effect and the absence of degradation cannot be attributed to changes in VHL ligase levels. The gradual or sudden onset of the hook effect suggests a differential interaction of SIM1 with BRD2 and BRD4 at higher concentrations. Imaide et al. demonstrated that SIM1 exhibits slightly different binding affinities for BRD2 compared to BRD4 using surface plasmon resonance (SPR) (fig. 5d), and our cellular thermal shift assay (CETSA)^5^ (Jafari et al. 2014) on the lysate of naïve HEK293 cells also showed that at high concentrations, SIM1 and MZ1 have a higher thermal stabilization with BRD2 compared to BRD4 (Fig. 2d).

## Discussion

The trivalent SIM1 degrader offers evidence that it is feasible to leverage avidity to develop highly efficacious degraders. However, even trivalent PROTACs can lead to hook effects, and it is vital to consider the context in which such degraders are evaluated in order to discern the contributions arising from increased avidity and those stemming from other factors (Fig. 2e). Factors such as target splice variants, the behavior of tagged versus non-tagged proteins, and differences between low and high efflux cells all require careful consideration.

## Figure Legends

**Supplementary Fig. 1. a**, Quantitative analysis of loss of luminescence due to degradation HiBiT-tagged BRD4 protein using 13-point dose response for LgBiT, HiBiT-BRD4 in HEK293 cells following 4 h treatment with MZ1, SIM1 or DMSO (3 technical replicates). **b**, Expression level of *ABCB1, ACTB, BRD4, PARK7, CRBN, KEAP1, SOD1, and VHL* in MV4-11, K562, NB4, HL60, U937, HEK293 (HEK) and HEL92.1.7 cells. Data extracted from Depmap portal. **c**, Immunoblot analysis of MDR1 protein level in HEL92.1.7, HL-60, K-562, MV4-11, NB-4, and U937 cells.

