## Supplementary figures and images for "Unveiling the Power of PROTAC Valency: Navigating Cell Type-Specific Hook Effects"

### Supplemental Figure 1

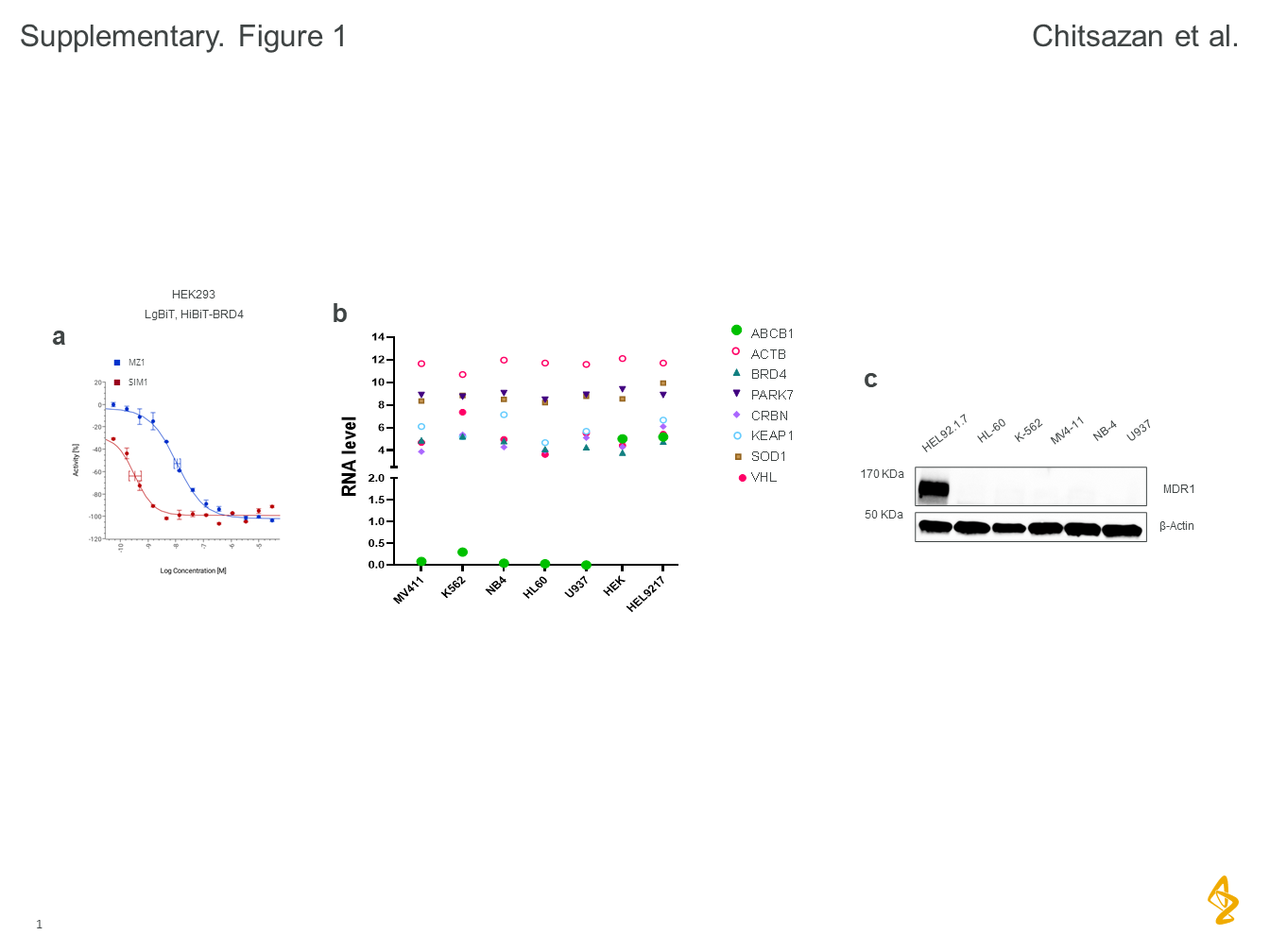
